# Cancer Predisposition Sequencing Reporter (CPSR): a flexible variant report engine for high-throughput germline screening in cancer

**DOI:** 10.1101/846089

**Authors:** Sigve Nakken, Vladislav Saveliev, Oliver Hofmann, Pål Møller, Ola Myklebost, Eivind Hovig

**Affiliations:** Department of Tumor Biology, Institute for Cancer Research, Oslo University Hospital, Norway; Centre for Cancer Cell Reprogramming, Institute of Clinical Medicine, Faculty of Medicine, University of Oslo, Norway; Centre for Cancer Research, University of Melbourne, Australia; Department of Clinical Science, University of Bergen, Norway; Western Norway Familial Cancer Center, Haukeland University Hospital, Bergen, Norway; Centre for Bioinformatics, Department of Informatics, University of Oslo, Norway

## Abstract

The value of high-throughput germline genetic testing is increasingly recognized in clinical cancer care. Disease-associated germline variants in cancer patients are important for risk management and surveillance, surgical decisions, and can also have major implications for treatment strategies since many are in DNA repair genes. With the increasing availability of high-throughput DNA sequencing in cancer clinics and research, there is thus a need to provide clinically oriented sequencing reports for germline variants and their potential therapeutic relevance on a per-patient basis. To meet this need we have developed the Cancer Predisposition Sequencing Reporter (CPSR), an open-source computational workflow that generates a structured report of germline variants identified in known cancer predisposition genes, highlighting markers of therapeutic, prognostic, and diagnostic relevance. A fully automated variant classification procedure based on more than 30 refined ACMG criteria represents an integral part of the workflow. Importantly, the set of cancer predisposition genes profiled in the report can be flexibly chosen from more than 40 virtual gene panels established by scientific experts, enabling customization of the report for different screening purposes and clinical contexts. The report can be configured to also list actionable secondary variant findings as recommended by ACMG, as well as the status of low-risk variants from genome-wide association studies in cancer. CPSR demonstrates superior sensitivity and comparable specificity for the detection of pathogenic variants when compared to existing algorithms. Technically, the tool is implemented in Python/R, and is freely available through Docker technology. Source code, documentation, example reports, and installation instructions are accessible via the project GitHub page: https://github.com/sigven/cpsr.

## 1 Introduction

A considerable fraction of human cancers is rooted in rare pathogenic germline mutations in cancer predisposition genes^1^. Screening of cancer patients for predisposing germline alterations may yield valuable decision support for risk-reducing interventions and surveillance, and has also proven its significance for the application of platinum-based chemotherapy and targeted drugs ^2,3^.

High-throughput screening for a broad collection of cancer predisposition genes is currently feasible due to technological advances in genome-wide DNA sequencing. The accuracy of variant detection algorithms has improved substantially, producing consistent and highly accurate results, particularly for single point mutations ^4^. On the other hand, the ability to interpret variant findings in terms of clinical significance and actionability still represents a major challenge. To our knowledge, no freely available bioinformatics tool aims to transform raw germline variant sets to structured and interactive reports for clinical interpretation on a per-patient basis. Efforts in this area have focused primarily on the implementation of algorithms for variant pathogenicity classification, which lies at the core of clinical variant interpretation. Multiple tools and algorithms for variant classification according to published guidelines by the American College of Medical Genetics and Genomics (ACMG) have been developed, the most relevant ones in the field of cancer being CharGer, SherLoc, and PathoMAN^5–7^. The comprehensive classification procedure outlined in Invitae’s SherLoc framework is however not available as open-source software, and the limited web-based service offered by PathoMAN is inconvenient for integration in high-throughput analysis environments. Furthermore, given the sensitive nature of DNA sequencing data from cancer patients, which is under strict regulations in most countries, it is frequently a necessity to choose stand-alone workflows over public web-based interpretation solutions. Also, neither of the above-mentioned tools and algorithms provide structured genome reports on a case-by-case basis, and where the report content can be customized according to the cancer condition in question. Summing up, although the generation of informative variant interpretation reports constitutes an essential output of high-throughput cancer sequencing workflows, there is currently a shortage of flexible solutions for this in the open-source software landscape.

Here, we present a flexible bioinformatics tool that generates personal genome reports in the context of cancer predisposition and inherited cancer syndromes. Cancer Predisposition Sequencing Reporter (CPSR) can be easily integrated with standard variant calling output from both whole-genome, exome or targeted gene panel sequencing, and produces structured and interactive variant reports that highlight findings with clinical implications.

## 2 Construction and Content

CPSR is implemented as a stand-alone bioinformatics workflow in Python and R. By design, it is therefore well suited for integration with workflows for high-throughput sequencing, as opposed to purely web-based solutions. Technically, CPSR builds upon our previously developed framework for the analysis of somatic mutations in tumor genomes, the Personal Cancer Genome Reporter (PCGR)^8^. To facilitate reproducibility and ease of use, the tool can be installed either as a Dockerized application or through a Conda package, the latter probably being the preferred choice in high-performance computing environments. In addition to the actual software and configuration files, users need to download a dedicated data bundle, which contains the underlying databases that CPSR is using for functional variant annotation and as a basis for classification and reporting. CPSR supports both of the recent assembly versions of the human genome (i.e. *grch37* and *grch38*). Installation instructions and other information regarding configuration, versions of software and underlying databases, and input/output files, are available from the project GitHub page (https://github.com/sigven/cpsr), and also through the CPSR documentation website (https://cpsr.readthedocs.io).

The input to CPSR is a single file with DNA variants (SNVs/InDels) detected from germline variant calling, encoded in the standard single-sample VCF format. CPSR automatically detects the genotype (homozygous/heterozygous) of input variants if these are formatted according to the correct standard in the VCF file. The workflow proceeds with four major steps, which are described in detail below (schematically illustrated in Figure 1).

**Figure 1:**
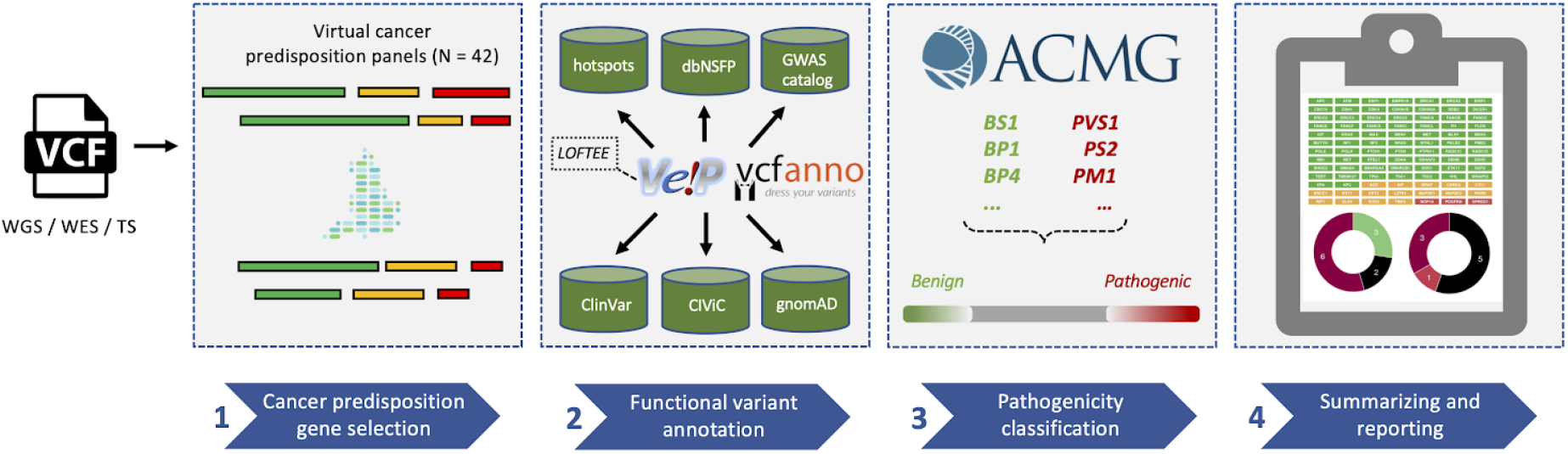
CPSR workflow with key databases and underlying software, illustrating how the query variant set from germline variant calling (formatted as VCF) is subject to four main steps for predisposition interpretation. Locus filtering against a selected cancer predisposition gene panel from the Genomics England PanelApp, where colors indicate confidence of association to phenotype, from diagnostic-grade in green to low-level confidence genes in red (**step 1**). Annotation through VEP and *vcfanno* with functional variant annotations: variant consequences by VEP, mutation hotspots from cancerhotspots.org, *in silico* deleteriousness predictions from dbNSFP, loss-of-function predictions through VEP’s LOFTEE plugin, population allele frequencies from gnomAD, germline biomarkers from CIViC, and low-risk alleles from NHGRI-EBI GWAS Catalog (**step 2**). Pathogenicity classification of novel variants according to a cancer-dedicated implementation of refined ACMG criteria (**step 3**). Aggregation and structuring of the results in a tiered cancer predisposition report (**step 4**). Abbreviations: VEP = Variant Effect Predictor; LOFTEE = Loss-Of-Function Transcript Effect Estimator; VCF = Variant Call Format; dbNSFP = database of non-synonymous functional predictions; CIViC = Clinical Interpretations of Variants in Cancer, ACMG = American College of Medical Genetics and Genomics; gnomAD = Genome Aggregation Database; WGS = Whole-Genome Sequencing; WES = Whole-Exome Sequencing; TS = Targeted Sequencing

### Selection of targets for reporting - virtual cancer predisposition gene panels

In order to serve a wide range of clinical cases, CPSR can produce variant reports that are dedicated towards predisposition genes for specific tumor types or cancer syndromes. In the initial step of the workflow, we exploit virtual gene panels as available from the Genomics England PanelApp, a crowdsourcing initiative in which scientific experts are evaluating risk genes for more than 40 different hereditary cancer conditions on a continuous basis^9^. Technically, variants in the input VCF file are filtered against the gene panel of choice (encoded as a BED file) to ensure that variants analyzed are restricted to the panel genes only. When selecting a panel from PanelApp for analysis in CPSR, the user may also restrict the analysis to genes with a high level of disease association only (i.e. diagnostic-grade or “GREEN” genes according to PanelApp nomenclature).

In addition to predefined panels from PanelApp, the user can choose to screen variants within a comprehensive exploratory panel intended for research use (i.e. a “superpanel”), containing cancer predisposition genes gathered from multiple sources. The superpanel includes a total of 335 protein-coding genes, containing all genes from PanelApp panels available in CPSR, genes curated in the Cancer Gene Census (COSMIC), those profiled in TCGA’s PanCancer analysis of germline variants, and other user-contributed genes deemed relevant for cancer predisposition. Importantly, users may also flexibly define their own virtual screening panel from the set of genes in the exploratory superpanel.

Information on dominant versus recessive inheritance patterns for the various inherited cancer syndromes is largely harvested from the Genomics England PanelApp, with some additions from two other large-scale sequencing studies of cancer genes^1,10^. Information related to the mechanism of disease (loss-of-function vs. gain-of-function) per gene has been collected from the study by Maxwell et al. ^10^. Disease-related gene properties are exploited during automated variant classification according to ACMG criteria (section *Automated variant pathogenicity classification* outlined below).

### Functional variant annotation

The second step of the workflow utilizes two open-source tools, Variant Effect Predictor (VEP) and *vcfanno*, to provide comprehensive functional annotations of all input variants ^11,12^.

Gene variant consequences are determined by VEP, using GENCODE as the gene and transcript reference model. Cross-references to RefSeq transcripts are provided in the output whenever this is available through Ensembl’s transcript database. Although a single variant frequently affects multiple transcripts in a given gene, CPSR reports a single main consequence per variant, using VEP’s internal ranking routine to pick the most important transcript-specific consequence, a ranking that can be configured by the user. Notably, variants with a putative loss-of-function consequence (i.e. stopgain, frameshift and splice site disruption), which are of major importance when it comes to pathogenic germline variants in cancer, are subject to careful evaluation and filtering through the LOFTEE plugin in VEP. Specifically, LOFTEE assigns confidence to a loss-of-function variant based on multiple features, such as transcript location, ancestral allele state, and intron size and donor site nature (for splice site mutations). The relative location of variants with respect to intron-exon borders are also derived from VEP’s output.

Through the use of *vcfanno*, the second workflow step will also annotate the input variants with data from multiple open-access variant datasets of relevance for cancer predisposition and functional variant effect (Figure 1). These datasets include information related to pre-classified variants in ClinVar (phenotypes, review status etc.), population-specific allele frequencies (gnomAD, non-cancer subset), known mutational hotspots in cancer (cancerhotspots.org), precomputed *insilico* deleteriousness predictions of missense and splice site variants (dbNSFP and dbSCSNV), low-risk risk alleles identified from genome-wide association studies of cancer phenotypes (GWAS catalog), and importantly, germline biomarkers of relevance for prognosis, diagnosis or therapeutic regimens retrieved from the Clinical Interpretations of Variants in Cancer resource (CIViC)^13–19^. Through annotations from CIViC, we can effectively show which germline variants in the query that, according to published evidence from clinical trials or case reports, are likely to have therapeutic implications. A prominent example relates to cases with increased sensitivity to poly(ADP-ribose) polymerase (PARP) inhibitors elicited by pathogenic variants in BRCA1/2 genes ^20^.

### Automated variant pathogenicity classification

The occurrence of rare variants that have not yet received any classification or interpretation (i.e. in ClinVar) is a common scenario in germline sequencing of cancer patients. To guide the interpretation of these variants, CPSR provides an automated pathogenicity classification in which the collection of variant annotations in step two (i.e. consequence type, predicted functional effect, and population frequencies), along with information on disease mechanism and mode of inheritance per cancer predisposition gene is exploited. Specifically, CPSR conducts a standard five-level (Benign/Likely Benign (B/LB), VUS, Likely Pathogenic/Pathogenic (P/LP)) variant pathogenicity classification^21^, serving similar functionality to the open-source tools offered by CharGer and PathoMAN.

The classification procedure employed by CPSR is built largely upon the foundations established by the SherLoc algorithm, which made substantial refinements to the original ACMG guidelines for variant classification^7^. In general, each ACMG criterion specifies particular properties of variants, such as population allele frequency and predicted functional effect, that supports a pathogenic or benign variant nature. Furthermore, in the approach proposed by SherLoc, each ACMG criterion is weighted with positive or negative point scores that reflect their relative strength of importance with respect to classification. For all criteria that match with a given variant, scores are ultimately aggregated to obtain a single variant pathogenicity score. Notably, the specific combinations of software (e.g. VEP and LOFTEE) and annotation databases (e.g. GENCODE, gnomAD) used in CPSR, as well as the customization of the criteria towards the disease phenotype (e.g. using cancer mutation hotspots to highlight important amino acids), are in effect providing a unique, cancer-dedicated variant classification procedure. The details of each ACMG evidence criterion implemented in CPSR, as well as their associated point scores, can be found in Supplementary Table 1.

Thresholds for converting variant pathogenicity scores to five-level classifications were calibrated through a comparison with existing ClinVar classifications (November 2020 release). In our calibration, we considered ClinVar variants in cancer predisposition genes (superpanel set, n = 335), limited to those with a review status of minimum two stars, the latter to minimize the impact of low-confident variant interpretations. The relationship between ClinVar classification status and variant pathogenicity scores calculated by CPSR is illustrated in Figure 2, and thresholds that were set ensured high concordance (agreement on 93.2% of all P/LP classified variants in ClinVar, 95.5% for VUS variants, and 96.4% for B/LB variants).

**Figure 2:**
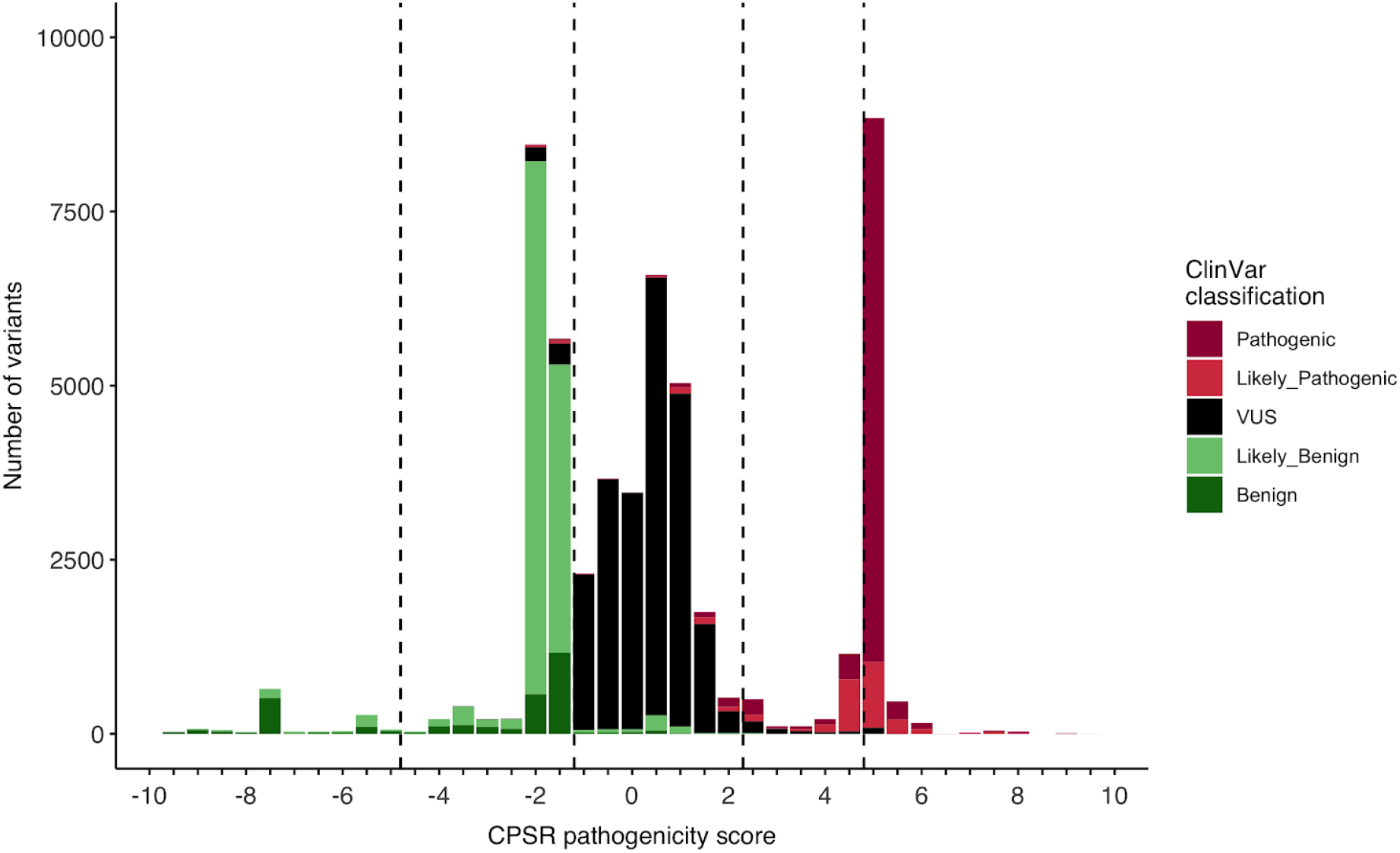
Calibration of CPSR pathogenicity score thresholds against ClinVar variants with a known classification (minimum two review stars). The complete distribution was calculated for variants in cancer predisposition genes (n = 335) and was used to determine suitable CPSR thresholds for P/LP/VUS/LB/B classifications, as indicated with the vertical dashed lines (Pathogenic: [5, ], Likely Pathogenic: [2.5, 4.5], VUS: [−1, 2.0], Likely Benign: [−4.5, −1.5], Benign: [, −5])

Finally, we compared the sensitivity and specificity of our classification algorithm with the two algorithms provided through CharGer (i.e. custom and ACMG-based). Here, we used an established benchmark set from the Pediatric Cancer Germline Project (PCGP) in which manual variant classifications defined by a panel of clinical geneticists constitute the gold standard (Supplementary Materials). For P/LP variants (n = 105), classification with CPSR achieved a sensitivity of 74.3%, which is higher than what was obtained with either of CharGer’s two algorithms (72.4% for the custom and 56.2% for the ACMG-based, respectively). Of all panel-determined non-pathogenic variants (n = 683), CPSR classified 16 variants as pathogenic, translating to a specificity of 97.7%, an intermediate of the rates produced by CharGer’s algorithms (98.1% for the ACMG-based and 97.3% for the custom, respectively).

### Variant report generation

The final step of the workflow exploits the R Markdown framework to display all variant findings in a structured and interactive variant report^22^. Of note is that additional output formats are also available to the user, i.e. annotated VCF, JSON, and TSV (tab-separated values). The TSV output can be utilized to collect results from multiple cases that have been analyzed with CPSR, which represents a common scenario in large research studies. An example HTML report can be downloaded for exploration here: https://doi.org/10.5281/zenodo.4050913.

The interactive HTML report is organized into four main sections: *Settings*, *Summary of Findings*, *Germline SNVs/InDels*, and *Documentation. Settings* indicate report and analysis configurations, as well as information regarding the virtual screening panel, while the *Documentation* section lists all versions of underlying databases and third-party software. These two sections thus serve to ensure reproducibility and transparency of the complete analysis workflow. *Summary of findings* provides the user with overall statistics with respect to classifications of variants found for the given case, both for variants already existing in ClinVar, and for novel variants without records in ClinVar.

The main content of the report is contained within the section named *Germline SNVs/InDels*, where details of all variants are structured in interactive tables, and where the user can explore and filter variant data for various types of annotations, e.g. population frequency, consequence type, or existing phenotype associations. For variants of uncertain significance (VUS), which frequently make up the largest group of variants, the report importantly enables the use of the CPSR pathogenicity score to prioritise potential borderline cases. A dedicated biomarker section lists input variants that can have therapeutic implications or otherwise influence prognosis or diagnosis, and also allows the user to investigate the supporting literature and evidence for such associations. Finally, the user can opt to list secondary pathogenic variants, as recommended by ACMG^23^, and also a potential overlap of input variants with low-to-moderate risk alleles found for cancer phenotypes in genome-wide association studies (GWAS).

## 3 Discussion

Knowledge on pathogenic variants in cancer-predisposing genes, and their relationships to systemic therapy choices, is emerging and evolving on a continuous basis. The quality and contents of the report produced with CPSR will thus advance accordingly, as underlying databases are updated. In particular, information regarding the mode of inheritance and the mechanism of action is currently not well established for a significant number of inherited cancers. Filling this gap is likely to improve variant classification in a number of genes.

The automated variant classification procedure implemented in CPSR demonstrated improved sensitivity over existing algorithms provided with CharGer. One should however note that simple comparisons of classification algorithms must be interpreted cautiously, primarily due to the fact that algorithms for variant classification are frequently configurable through a multitude of parameters. It should also be emphasized that automated procedures are intended primarily to guide the classification, and where borderline cases either way should be manually reviewed.

We acknowledge that additional datasets and analyses can add useful extra dimensions to the cancer predisposition report. A future version of CPSR should accept germline DNA copy number variants as an additional input type. Important pharma- and radiogenomic risk variants may further be incorporated during reporting, as well as a framework for calculation of polygenic risk scores ^24^.

## 4 Conclusion

Evidence-based personal cancer treatment based on genetic testing is an important goal in oncology. CPSR provides a documented tool to reach this goal.

## Acknowledgements

This work was supported by grant #221580 from the Norwegian Research Council to the Norwegian Cancer Genomics Consortium.

## Data Availability Statement

Source code and documentation of CPSR can be found through the project GitHub repository (https://github.com/sigven/cpsr). Supplementary materials, including supplementary tables, scripts and datasets for calibration/benchmark calculations are available here: https://doi.org/10.5281/zenodo.4291042

## References

1. Huang K-L, Mashl RJ, Wu Y, Ritter DI, Wang J, Oh C, Paczkowska M, Reynolds S, Wyczalkowski MA, Oak N, Scott AD, Krassowski M, et al. Pathogenic Germline Variants in 10,389 Adult Cancers. Cell 2018;173:355–70.e14.

2. Thavaneswaran S, Rath E, Tucker K, Joshua AM, Hess D, Pinese M, Ballinger ML, Thomas DM. Therapeutic implications of germline genetic findings in cancer. Nat Rev Clin Oncol 2019;16:386–96.

3. Lincoln SE, Nussbaum RL, Kurian AW, Nielsen SM, Das K, Michalski S, Yang S, Ngo N, Blanco A, Esplin ED. Yield and Utility of Germline Testing Following Tumor Sequencing in Patients With Cancer. JAMA Netw Open 2020;3:e2019452.

4. Chen J, Li X, Zhong H, Meng Y, Du H. Systematic comparison of germline variant calling pipelines cross multiple next-generation sequencers. Sci Rep 2019;9:9345.

5. Scott AD, Huang K-L, Weerasinghe A, Mashl RJ, Gao Q, Martins Rodrigues F, Wyczalkowski MA, Ding L. CharGer: clinical Characterization of Germline variants. Bioinformatics 2019;35:865–7.

6. Ravichandran V, Shameer Z, Kemel Y, Walsh M, Cadoo K, Lipkin S, Mandelker D, Zhang L, Stadler Z, Robson M, Offit K, Vijai J. Toward automation of germline variant curation in clinical cancer genetics. Genet Med 2019;21:2116–25.

7. Nykamp K, Anderson M, Powers M, Garcia J, Herrera B, Ho Y-Y, Kobayashi Y, Patil N, Thusberg J, Westbrook M, The Invitae Clinical Genomics Group, Topper S. Sherloc: a comprehensive refinement of the ACMG–AMP variant classification criteria. Genet Med 2017;19:1105.

8. Nakken S, Fournous G, Vodák D, Aasheim LB, Myklebost O, Hovig E. Personal Cancer Genome Reporter: variant interpretation report for precision oncology. Bioinformatics 2018;34:1778–80.

9. Martin AR, Williams E, Foulger RE, Leigh S, Daugherty LC, Niblock O, Leong IUS, Smith KR, Gerasimenko O, Haraldsdottir E, Thomas E, Scott RH, et al. PanelApp crowdsources expert knowledge to establish consensus diagnostic gene panels. Nat Genet 2019;51:1560–5.

10. Maxwell KN, Hart SN, Vijai J, Schrader KA, Slavin TP, Thomas T, Wubbenhorst B, Ravichandran V, Moore RM, Hu C, Guidugli L, Wenz B, et al. Evaluation of ACMG-Guideline-Based Variant Classification of Cancer Susceptibility and Non-Cancer-Associated Genes in Families Affected by Breast Cancer. Am J Hum Genet 2016;98:801–17.

11. McLaren W, Gil L, Hunt SE, Riat HS, Ritchie GRS, Thormann A, Flicek P, Cunningham F. The Ensembl Variant Effect Predictor. Genome Biol 2016;17:122.

12. Pedersen BS, Layer RM, Quinlan AR. Vcfanno: fast, flexible annotation of genetic variants. Genome Biol 2016;17:118.

13. Griffith M, Spies NC, Krysiak K, McMichael JF, Coffman AC, Danos AM, Ainscough BJ, Ramirez CA, Rieke DT, Kujan L, Barnell EK, Wagner AH, et al. CIViC is a community knowledgebase for expert crowdsourcing the clinical interpretation of variants in cancer. Nat Genet 2017;49:170–4.

14. Liu X, Jian X, Boerwinkle E. dbNSFP: a lightweight database of human nonsynonymous SNPs and their functional predictions. Hum Mutat 2011;32:894–9.

15. Landrum MJ, Lee JM, Riley GR, Jang W, Rubinstein WS, Church DM, Maglott DR. ClinVar: public archive of relationships among sequence variation and human phenotype. Nucleic Acids Res 2014;42:D980–5.

16. Buniello A, MacArthur JAL, Cerezo M, Harris LW, Hayhurst J, Malangone C, McMahon A, Morales J, Mountjoy E, Sollis E, Suveges D, Vrousgou O, et al. The NHGRI-EBI GWAS Catalog of published genome-wide association studies, targeted arrays and summary statistics 2019. Nucleic Acids Res 2019;47:D1005–12.

17. Chang MT, Shrestha Bhattarai T, Schram AM, Bielski CM, Donoghue MTA, Jonsson P, Chakravarty D, Phillips S, Kandoth C, Penson A, Gorelick A, Shamu T, et al. Accelerating discovery of functional mutant alleles in cancer. Cancer Discov [Internet] 2017;Available from: http://dx.doi.org/10.1158/2159-8290.CD-17-0321

18. Karczewski KJ, Francioli LC, Tiao G, Cummings BB, Alföldi J, Wang Q, Collins RL, Laricchia KM, Ganna A, Birnbaum DP, Gauthier LD, Brand H, et al. The mutational constraint spectrum quantified from variation in 141,456 humans. Nature 2020;581:434–43.

19. Jian X, Boerwinkle E, Liu X. In silico prediction of splice-altering single nucleotide variants in the human genome. Nucleic Acids Res 2014;42:13534–44.

20. Fong PC, Boss DS, Yap TA, Tutt A, Wu P, Mergui-Roelvink M, Mortimer P, Swaisland H, Lau A, O’Connor MJ, Ashworth A, Carmichael J, et al. Inhibition of poly(ADP-ribose) polymerase in tumors from BRCA mutation carriers. N Engl J Med 2009;361:123–34.

21. Plon SE, Eccles DM, Easton D, Foulkes WD, Genuardi M, Greenblatt MS, Hogervorst FBL, Hoogerbrugge N, Spurdle AB, Tavtigian SV, IARC Unclassified Genetic Variants Working Group. Sequence variant classification and reporting: recommendations for improving the interpretation of cancer susceptibility genetic test results. Hum Mutat 2008;29:1282–91.

22. Allaire JJ, Yihui Xie MJ, Javier L, Kevin U, Aron A, Hadley W, Joe C, Winston C, Richard I. Rmarkdown: Dynamic Documents for R [Internet]. 2019. Available from: https://github.com/rstudio/rmarkdown

23. Kalia SS, Adelman K, Bale SJ, Chung WK, Eng C, Evans JP, Herman GE, Hufnagel SB, Klein TE, Korf BR, McKelvey KD, Ormond KE, et al. Recommendations for reporting of secondary findings in clinical exome and genome sequencing, 2016 update (ACMG SF v2.0): a policy statement of the American College of Medical Genetics and Genomics. Genet Med 2017;19:249–55.

24. Torkamani A, Wineinger NE, Topol EJ. The personal and clinical utility of polygenic risk scores. Nat Rev Genet 2018;19:581–90.

