## Supplementary Materials for "Cancer Predisposition Sequencing Reporter (CPSR): a flexible variant report engine for high-throughput germline screening in cancer"

11/25/2020

### CPSR variant classification

#### Calibration

In order to establish thresholds for conversion of CPSR pathogenicity scores to categorical variant classifications, we considered high-confident variant classifications from ClinVar. Specifically, we downloaded the ClinVar VCF file from the ClinVar FTP site (clinvar_20201031.vcf.gz), build grch37, and analyzed variants with a review status of minimum two stars with CPSR for all predisposition genes (CPSR panel 0, n = 335 genes). Through an iterated procedure, we determined thresholds for the pathogenicity score that provided an optimal separation of P/LP, VUS, and B/LB variants. The table below indicates how many ClinVar variants were present per classification category, and how many were correctly classified by CPSR based the chosen thresholds (outlined in Figure 2 of main manuscript). Scripts and datasets for the calibration calculations are available here: https://doi.org/10.5281/zenodo.4290411

| Classification | Variants - ClinVar | Variants - CPSR | Agreement (%) |
|---|---|---|---|
| PLP | 12354 | 11511 | 93.18 |
| VUS | 23638 | 22584 | 95.54 |
| BLB | 17296 | 16679 | 96.43 |

#### Benchmark

To assess the sensitivity and specificity of CPSR's pathogenic variant classification, we used a dataset of 788 unique variants from the Pediatric Cancer Germline Project (PCGP) in which classifications had been manually established by a panel of clinical geneticists (Zhang et al. 2015). This is the same dataset that was used to benchmark the algorithms (ACMG and custom) provided with CharGer (Scott et al. 2019). Note that our benchmark results are computed based on the set of unique PCGP variants only, as opposed to CharGer, which considered all case variants in PCGP, effectively considering the same variant multiple times when it occurs in more than one case. We classified the variant set (n = 788) with CPSR, and compared the classifications against PCGP's gold standard classifications (results presented in main manuscript). The complete benchmark variant set with all classifications can be found in Supplementary Table 2. Scripts and datasets for the benchmarking procedure are available here: https://doi.org/10.5281/zenodo.4290411

### Supplementary Tables (.xlsx)

**Table S1** - Table with detailed descriptions of all ACMG evidence criteria and corresponding scores that collectively form the basis for variant classification in CPSR. Criteria that support a pathogenic effect are highlighted in red, while criteria that support a benign effect are highlighted in green.

**Table S2** - Table with complete benchmark variant dataset from PCGP with classifications from CPSR, CharGer (ACMG), and CharGer (Custom)

### References

Scott, Adam D, Kuan-Lin Huang, Amila Weerasinghe, R Jay Mashl, Qingsong Gao, Fernanda Martins Rodrigues, Matthew A Wyczalkowski, and Li Ding. 2019. "CharGer: Clinical Characterization of Germline Variants." *Bioinformatics* 35 (5): 865–67. http://dx.doi.org/10.1093/bioinformatics/bty649.

Zhang, Jinghui, Michael F Walsh, Gang Wu, Michael N Edmonson, Tanja A Gruber, John Easton, Dale Hedges, et al. 2015. "Germline Mutations in Predisposition Genes in Pediatric Cancer." *N. Engl. J. Med.* 373 (24): 2336–46. http://dx.doi.org/10.1056/NEJMoa1508054.
